# Mixed-cropping systems of different rice cultivars have grain yield and quality advantages over mono-cropping systems

**DOI:** 10.1101/317545

**Authors:** Meijuan Li, Jiaen Zhang, Shiwei Liu, Umair Ashraf, Shuqing Qiu

## Abstract

Mixed-cropping system is a centuries-old cropping technique that is still widely practiced in the farmers’ field over the globe. Increased plant diversity enhances farmland biodiversity, which would improve grain yield and quality; however, the impacts of growing different rice cultivars simultaneously were rarely investigated. In present study, five popular rice cultivars were selected and ten mixture combinations were made according to the growth period, plant height, grain yield and quality, and pest and disease resistance. Seedlings of the five cultivars and ten mixture combinations (mixed-sowing of the seeds in an equal ratio, then mixed-transplanting and finally mixed-harvesting) were grown in plastic pots under greenhouse during the early and late growing seasons in 2016. Results showed that, compared with the corresponding mono-cropping systems, almost all combinations of the mixed-cropping systems have advantages in yield related traits and grain quality. Compared with the mono-cropping systems in the early and late growing seasons in 2016, mixed-cropping systems increased the number of spikelets per panicle, seed-setting rate, and grain weight per pot and harvest index by 19.52% and 5.77%, 8.53% and 4.41%, 8.31% and 4.61%, and 10.26% and 6.98%, respectively (paired t-test). In addition, mixed-cropping systems reduced chalky rice rate and chalkiness degree by 33.12% and 43.42% and by 30.11% and 48.13% in the early and late growing seasons, respectively (paired t-test). These results may be due to enhanced SPAD indexes and photosynthetic rates at physiology maturity in mixed-cropping systems. In general, it was found that mixed-cropping with different rice cultivars have potential for increasing grain yield and improving grain quality.

## Introduction

Rice (*Oryza sativa* L.) is the primary food source for more than one-third of the world’s population [1]. The socio-economic and climatic factors are negatively affecting rice production worldwide [2–4]. Recently, efforts have been made to modernize the rice production systems by replacing traditional farming systems with improved production practices and agricultural mechanization [5]. The traditional farming systems at small and large scale have greatly been replaced with intensified and highly mechanized mono-culture based cropping systems [6–8]. These production systems are largely rely on chemical fertilizers and pesticides which may cause significant surface and groundwater contamination and potential health risks [9,10]. The intensive cropping increases the chances of soil erosion, greenhouse gas emission, pest resistance, and the loss of biodiversity. Hence, sustainable crop production systems are essential over the long term to meet the consumers’ demand for better-quality food products [11,12].

Multi-cropping, refers to as ‘intercropping’ or ‘mixed-cropping’, is the growing multiple crop species/cultivars simultaneously in the same field for a significant part of their life cycle [6,13]. Numerous studies have reported how ecological processes result in yield advantages in mixed-cropping systems compared to those in mono-cropping systems. For example, studies on grasslands have shown that multi-species produced 15% higher yields than mono-crops [14]. Mixed-cropping has also been shown to produce 1.7 times more biomass than single species mono-cropping and to be 79% more productive than mono-cropping system [15]. In addition to yield, there are many other benefits of multiple-cropping such as enhanced soil fertility by intercropping with nitrogen-fixers [16], increased resilience against pests and diseases [17], and increased abiotic stress tolerance have also been previously reported [18,19]. These effects are attributed to higher levels of genetic diversity within those systems [6,7,16,20,21].

Furthermore, mixed-cropping systems are mainly used in tropical, small-scale subsistence farming [6], however, it may have some practical issues, such as drilling, sowing, spraying and harvesting practices etc. Differing growth cycles and requirements for nutrients and pesticides make it difficult for growers to adapt new systems to manage and harvest mixed crops [6,21,22].

However, it is easy to operate the mixed-cropping systems when several cultivars belong to the same species(eco-types), are mixed and seeded in the same field. Meanwhile, mixed-cropping systems may have higher resistance to diseases or pests, higher production, better grain quality, and may allow the situ production of formula rice (products with different rice varieties) easily. Till now, no work has been done to examine the effects of cultivating different rice genotypes in different combination on grain yield and quality of rice. Therefore, in this study, the seeds of different rice cultivars (possess different growth periods, plant heights, disease and pest resistance, grain yield and quality), were mixed and planted. It was hypothesized that a mixed-cropping system with different rice cultivars would have advantages over mono-cropping systems for a series of traits related to grain production and quality.

## Materials and Methods

Experiments were conducted in 2016 during the early (March-July) and late growing seasons (August-November) of rice in the greenhouse at the Eco–Farm in the campus of South China Agricultural University (113°21′E, 23°09′N), Guangzhou, China. This region has a humid subtropical monsoonal climate characterized by warm winter and hot summers with an average annual temperature between 20°C and 22°C.

### Experimental details

Five conventional indica rice cultivars of which four i.e., Yuenongsimiao, Yuxiangyouzhan, Huangguangyouzhan and Huanghuazhan were obtained from the Rice Research Institute of the Guangdong Academy of Agricultural Sciences and one i.e., Huahang 31 from the National Engineering Research Centre of Plant Breeding at South China Agricultural University (named as A, B, C, D and E, respectively) were used as plant material. Out of 26 possible 1:1 mixture combinations of five genotypes, we selected ten combinations i.e., BC, BE, AB, BCE, ACD, BDE, ABD, BCDE, ABDE and ABCDE according to the growth period, plant height, grain yield, grain qualities, and resistance to diseases and pests of the five cultivars. The seed mixtures (mixing the seeds in an equal ratio) were sown in PVC trays and then the 35-day-old seedlings were transferred to the soil containing plastic pots (60×45×29 cm) filled with 30 kg of soil from paddy fields and grown in the greenhouse during the early and late growing seasons in 2016. The five cultivars were also grown in pure stands in mono-cropping systems as the corresponding controls. The experimental soil was sandy loam containing 25.44 g kg−1 organic matter, 1.14 g kg−1 total nitrogen, 0.84 g kg−1 total phosphorous, 22.36 g kg−1 total potassium and had a pH of 5.98. Before transplanting, 100 g of organic fertilizer (N, P_2_O_5_, K_2_O ≥6%, organic matter ≥46%) was applied per pot. There were 3 replications for 5 cultivars and 10 mixtures whilst 6 hills (6 seedlings for each hill) with a planting space of 20 × 15 cm for each pot were maintained. A water layer of about 2-3 cm was maintained for the whole growth period.

### Trait measurements

The photosynthetic rate (Pn), relative chlorophyll contents (SPAD) and total aboveground dry weight (DW) were measured at physiological maturity. The maximum CO_2_ assimilation rate per unit area (Pn; μmol m-2 s-1) was measured between 9:00 and 11:00 am using a Li-6400 portable photosynthetic system (Li-6400, Li-Cor, USA). Based on preliminary trials, the photosynthetic photon flux density was set at 1000 μmol m-2 s-1 for all rice cultivars. Both of the ambient CO_2_ and air temperature were maintained at 390 μmol mol-1 and 28°C, respectively. Relative chlorophyll contents of flag leaf were estimated with a SPAD meter (SPAD-502, Osaka, Japan). The plants were kept in oven at 80°C till constant weight for determination of dry biomass.

Grain yield and its components were measured according to the methods described by Peng et al. (2004) [23]. At maturity stage, plants were sampled from the plastic pots and the panicles were cut off into straw and panicle individuals. All spikelets were separated from the rachis (by manual threshing), and divided into filled and unfilled rice by the Seeds Winnowing machine (CFY-II, 3.8m3 min, max air pressure 1300 pa, Hangzhou, China). The total number of spikelets, filled and unfilled, all of the half-filled spikelets were classifically taken and averaged. The number of spikelets per panicle, grain-filling percentage (100 × the number of filled spikelets / the total number of spikelets), and 1000-grain-weight were also calculated from sampled plants and averaged. Grain yield was recorded from each pot and the grain moisture was reduced to 14% by sun drying before being weighed.

Representative samples of about 250g of filled grains collected from each mono-cropping and mixed-cropping treatment were analyzed for grain quality. After dehulling and polishing rough rice, head rice (with length ≥ 3/4 of its total grain length) was weighed and used to calculate head rice yield. Physical traits such as chalky rice rate, chalkiness degree, grain length and width were scanned by a Plant Mirror Image Analysis (MICROTEK ScanMaker i800plus, Shanghai, China), and the image was processed with SC-E software (Hangzhou Wanshen Detection Technology Co., Ltd., Hangzhou, China). The standard iodine colorimetric method described in GB/T 15683-2008 (National Standard of the People’s Republic of China, 2008) was used to measure amylose content and the Coomassie Brilliant Blue Staining method was used to measure the grain soluble protein contents.

### Data analysis

The independent t-test was performed to evaluate the differences in traits between the mono-cropping and mixed-cropping treatments. For example, the data from A and B mono-cropping systems were pooled, and then compared with these data from AB mixed-cropping system. To assess the total effect of mixed-cropping, a paired analysis (t-test when data met assumptions of normality) and wilcoxon signed-rank test (when data did not meet the assumptions of normality) was carried out for all combinations and their mid-component average (the average of mixture components grown as mono-cropping). All the analyses were conducted in R 3.20 (R Foundation for Statistical Computing).

## Results

### Grain yield and its components

There were some significant differences in both yield and yield components between the mono-cropping and mixed-cropping systems for both early and late growing season in 2016 (Paired t-test, Tables 1 and 2). The spikelet per panicle, seed-setting rate, and grain weight per pot and harvest index were 19.52% and 5.77%, 8.53% and 4.41%, 8.31% and 4.61%, and 10.26% and 6.98% higher in the mixed-cropping systems than those in the mono-cropping systems in the early and late growing seasons, respectively. In the early growing season, the mixed-cropping system improved the number of spikelets per panicle and seed-setting rate, compared with mono-cropping system, but statistically non-significant (*P>0.05*) for some cases (Independent t-test). The grain weight per pot, only in the mixed-cropping systems of combinations of BC, ACD, BDE and ABDE, was significantly higher than that in the corresponding mono-cropping systems in the early growing season. Moreover, the grain weight was almost higher in the mixed-cropping systems in the early growing season and in most mixed-cropping combinations (except BE, ABD and ABCDE) in the late growing season, compared with the mono-cropping system. However, enhanced 1000-grain-weight was found only in some mixed-cropping systems in both the early and late growing seasons.

**Table 1.** Differences of yield and its components between mono-cropping and corresponding mixed-cropping systems with several cultivars in the early growing season of 2016. Note: Mean values (± SEs) of traits measured for mono-cropping systems and mixed-cropping systems. For each pair of combination we performed an independent t-test, and mean differences between mono-cropping systems and mixed-cropping systems were tested with paired t-tests. Mono, mono-cropping system; mix, mixed-cropping system. ^*^ *P*<0.05; ^**^ *P*<0.01.

| Treatments | The number of spikelet per panicle | Seed-setting rate (%) | Grain weight (g per pot) | 1000-grain -weight(g) | Harvest index |
| --- | --- | --- | --- | --- | --- |
| Independent t-test |  |  |  |  |  |
| Mono-AB | 46.05±5.72 | 63.83±1.88 | 52.13±3.63 | 20.11±0.43 | 0.39±0.02 |
| Mixed-AB | 56.05±1.25 | 70.14±0.29 | 53.76±3.41 | 18.61±1.12 | 0.45±0.00 |
| Mono-BC | 50.20±7.39 | 62.50±1.44 | 51.64±3.30 | 19.37±0.13 | 0.42±0.01 |
| Mixed-BC | 60.46±2.53 | 68.09±0.90* | 61.09±1.92 | 20.70±0.45** | 0.47±0.02* |
| Mono-BE | 53.11±8.78 | 65.79±2.76 | 49.65±2.41 | 18.86±0.36 | 0.38±0.02 |
| Mixed-BE | 58.73±0.69 | 68.66±2.32 | 49.63±1.10 | 17.80±1.01 | 0.37±0.02 |
| Mono-ABD | 43.09±4.01 | 66.48±1.81 | 52.97±2.38 | 20.39±0.32 | 0.40±0.01 |
| Mixed-ABD | 66.24±1.67** | 68.92±1.33 | 56.89±1.95 | 18.91±1.02 | 0.39±0.01 |
| Mono-ACD | 53.75±4.41 | 68.34±1.09 | 57.82±1.07 | 20.38±0.32 | 0.39±0.01 |
| Mixed-ACD | 64.21±1.79 | 77.63±1.84** | 62.47±0.61* | 23.25±0.61** | 0.46±0.01* |
| Mono-BCE | 57.46±6.08 | 65.62±1.84 | 52.74±2.21 | 19.02±0.26 | 0.39±0.01 |
| Mixed-BCE | 66.23±1.92 | 73.19±1.56* | 53.56±2.24 | 18.47±0.40 | 0.42±0.00 |
| Mono-BDE | 47.80±6.29 | 67.78±2.06 | 51.32±1.77 | 19.56±0.42 | 0.40±0.01 |
| Mixed-BDE | 65.19±1.55 | 74.81±1.20 | 63.49±1.33** | 20.93±0.25 | 0.46±0.01* |
| Mono-ABDE | 50.32±4.85 | 67.82±1.53 | 53.46±1.79 | 19.88±0.38 | 0.38±0.01 |
| Mixed-ABDE | 57.58±3.06 | 77.56±0.59** | 62.60±0.94* | 21.23±0.06 | 0.43±0.03 |
| Mono-BCDE | 52.39±5.23 | 67.16±1.59 | 53.22±1.65 | 19.51±0.32* | 0.40±0.01 |
| Mixed-BCDE | 57.04±1.81 | 70.40±1.40 | 53.64±1.66 | 17.94±0.60 | 0.39±0.00 |
| Mono-ABCDE | 53.49±4.21 | 67.32±1.27 | 54.55±1.54 | 19.78±0.31** | 0.39±0.01 |
| Mixed-ABCDE | 55.11±1.17 | 69.72±2.16 | 56.38±0.67 | 17.49±0.20 | 0.45±0.01* |
| Paired t-test |  |  |  |  |  |
| Mono-cropping | 50.77±1.33 | 66.26±0.59 | 52.95±0.69 | 19.68±0.17 | 0.39±0.00 |
| Mixed-cropping | 60.68±1.39** | 71.91±1.15** | 57.35±1.52* | 19.53±0.60 | 0.43±0.01** |

**Table 2.** Differences of yield and its components between mono-cropping and corresponding mixed-cropping systems with several cultivars in the late growing season of 2016. Note: Table explanations are provided in Table 1.

| Treatments | The number<br>of spikelet<br>per panicle | Seed-setting<br>rate (%) | Grain weight (g<br>per pot) | 1000-grain<br>-weight(g) | Harvest index |
| --- | --- | --- | --- | --- | --- |
| Independent t-test |  |  |  |  |  |
| Mono-AB | 60.26±3.39 | 72.47±0.95 | 62.56±1.00 | 21.41±0.65 | 0.44±0.00 |
| Mixed-AB | 63.06±1.47 | 74.81±0.81 | 65.60±0.12 | 22.16±0.92 | 0.46±0.01* |
| Mono-BC | 60.69±3.60 | 72.09±0.83 | 62.61±0.92 | 21.36±0.64 | 0.44±0.01 |
| Mixed-BC | 65.18±0.72 | 79.98±1.10** | 72.43±1.62** | 22.55±0.30 | 0.46±0.01* |
| Mono-BE | 61.17±3.76 | 73.33±1.33 | 61.71±1.84 | 21.07±0.73 | 0.42±0.02 |
| Mixed-BE | 66.93±1.20 | 72.41±0.99 | 60.55±2.18 | 22.18±0.74 | 0.44±0.01 |
| Mono-ABD | 59.71±2.25 | 73.00±0.69 | 63.04±0.73 | 21.64±0.46 | 0.43±0.00 |
| Mixed-ABD | 66.17±1.88 | 76.32±0.81* | 62.37±1.07 | 20.80±0.34 | 0.45±0.01 |
| Mono-ACD | 64.86±1.75 | 74.06±0.36 | 63.12±0.55 | 22.18±0.22 | 0.44±0.01 |
| Mixed-ACD | 63.57±0.55 | 76.19±0.54* | 66.73±1.10** | 22.68±0.38 | 0.47±0.00* |
| Mono-BCE | 63.58±2.75 | 73.45±0.89 | 62.05±1.23 | 21.43±0.52 | 0.43±0.01 |
| Mixed-BCE | 63.43±0.37 | 76.77±0.49 | 65.67±0.80 | 23.39±0.69 | 0.45±0.01 |
| Mono-BDE | 60.32±2.51 | 73.58±0.89 | 62.47±1.26 | 21.42±0.53 | 0.42±0.01 |
| Mixed-BDE | 68.73±1.77 | 79.10±1.94* | 67.29±1.01 | 23.69±0.68* | 0.52±0.03** |
| Mono-ABDE | 62.12±2.10 | 73.79±0.68 | 62.51±0.97 | 21.63±0.41 | 0.42±0.01 |
| Mixed-ABDE | 68.60±0.68 | 78.69±1.54** | 67.83±0.97* | 21.27±0.19 | 0.49±0.01** |
| Mono-BCDE | 62.34±2.16 | 73.60±0.68 | 62.53±0.95 | 21.60±0.41 | 0.42±0.01 |
| Mixed-BCDE | 66.34±0.95 | 78.53±1.37** | 64.32±0.80 | 22.98±0.23 | 0.45±0.00 |
| Mono-ABCDE | 63.38±1.82 | 73.77±0.55 | 62.56±0.79 | 21.73±0.34 | 0.43±0.01 |
| Mixed-ABCDE | 62.14±1.03 | 72.64±0.96 | 61.25±0.77 | 21.30±0.70 | 0.42±0.01 |
| Paired t-test |  |  |  |  |  |
| Mono-cropping | 61.84±0.54 | 73.31±0.20 | 62.52±0.13 | 21.55±0.09 | 0.43±0.00 |
| Mixed-cropping | 65.41±0.73** | 76.54±0.83** | 65.40±1.12* | 22.30±0.30* | 0.46±0.01** |

### Grain quality

For grain quality, chalky rice rate and chalkiness degree were significantly (*P*<0.05) decreased in the mixed-cropping systems in the early growing season, while other traits were not significantly different from those in the mono-cropping systems. Further, the mixed-cropping systems reduced the chalky rice rate and chalkiness degree by 33.12% and 43.42%, and 30.11% and 48.13% in the early and late growing seasons, respectively (Paired t-test, Tables 3 and 4). In the early season, only for several cases, grain milling and appearance, cooking and nutritional qualities of the mixed-cropping systems were significantly (*P*< 0.05) higher than those of the mono-cropping systems, whereas significant differences were found between mono-cropping and mixed-cropping systems in the late growing season (Independent t-test).

**Table 3.** Differences of grain quality between mono-cropping and corresponding mixed-cropping systems with several cultivars in the early growing season of 2016. Note: Table explanations are provided in Table 1.

| Treatment | Brown rice<br>rate (%) | Milled rice<br>rate (%) | Whole milled<br>rice rate<br>(%) | Amylose<br>content (%) | Protein<br>content (%) | Length/width | Chalky rice rate<br>(%) | Chalkiness<br>degree (%) |
| --- | --- | --- | --- | --- | --- | --- | --- | --- |
| Independent t-test |  |  |  |  |  |  |  |  |
| Mono-AB | 80.04±0.34 | 69.87±0.92 | 64.79±1.56 | 22.97±1.52 | 8.55±0.26 | 2.91±0.03* | 21.18±2.81 | 7.07±0.48* |
| Mixed-AB | 81.38±0.76 | 71.16±1.71 | 63.28±1.05 | 23.70±0.47 | 8.57±0.64 | 2.75±0.01 | 12.35±0.55 | 4.84±0.48 |
| Mono-BC | 80.35±0.41 | 69.91±0.93 | 63.37±1.04 | 21.68±2.16 | 8.57±0.31 | 2.86±0.04 | 22.18±2.43* | 6.75±0.60 |
| Mixed-BC | 80.88±0.43 | 70.34±1.16 | 68.21±1.45* | 22.47±0.29 | 9.35±0.48 | 2.90±0.07 | 12.44±0.82 | 5.02±0.52 |
| Mono-BE | 79.83±0.53 | 67.91±1.14 | 59.98±1.63 | 23.22±1.60 | 8.38±0.30 | 2.87±0.04 | 21.05±2.85 | 6.99±0.51* |
| Mixed-BE | 80.39±0.82 | 68.47±0.71 | 60.96±1.97 | 23.99±0.76 | 7.81±0.24 | 2.78±0.07 | 13.30±0.55 | 4.85±0.11 |
| Mono-ABD | 79.72±0.30 | 69.77±0.61 | 64.70±1.01 | 21.97±1.26 | 8.72±0.21 | 2.92±0.02 | 18.01±2.41 | 6.40±0.47 |
| Mixed-ABD | 78.84±0.82 | 69.89±0.96 | 61.75±0.57 | 19.89±1.06 | 8.37±0.83 | 2.94±0.08 | 12.29±0.26 | 4.87±0.17 |
| Mono-ACD | 79.68±0.30 | 70.28±0.36 | 65.99±0.69 | 18.94±0.89 | 8.73±0.19 | 2.90±0.03 | 14.56±0.94** | 5.55±0.20** |
| Mixed-ACD | 82.2±0.17** | 71.25±1.74 | 69.36±1.59* | 20.75±0.83 | 10.06±0.18** | 3.02±0.05 | 6.67±0.81 | 1.50±0.70 |
| Mono-BCE | 79.99±0.40 | 68.83±0.89 | 61.76±1.41 | 21.19±1.51 | 8.45±0.24 | 2.86±0.03 | 19.70±2.01 | 6.49±0.42* |
| Mixed-BCE | 78.43±0.48 | 67.53±0.12 | 64.76±0.53 | 18.39±0.69 | 9.74±0.15* | 2.77±0.05 | 13.93±1.14 | 4.45±0.35 |
| Mono-BDE | 79.57±0.39 | 68.46±0.80 | 61.49±1.30 | 22.14±1.31 | 8.61±0.24 | 2.89±0.03 | 17.92±2.42 | 6.35±0.47 |
| Mixed-BDE | 80.52±1.07 | 70.56±1.79 | 68.05±1.38* | 19.49±0.42 | 9.07±0.32 | 3.03±0.05* | 10.71±0.30 | 4.61±0.21 |
| Mono-ABDE | 79.60±0.29 | 69.00±0.67 | 63.16±1.30 | 21.53±1.03 | 8.59±0.19 | 2.90±0.02 | 17.19±1.84 | 6.29±0.35 |
| Mixed-ABDE | 80.48±0.32 | 69.94±1.47 | 67.48±1.22 | 21.65±0.87 | 9.13±0.34 | 2.96±0.05 | 12.33±0.43 | 5.04±0.08 |
| Mono-BCDE | 79.76±0.33 | 69.02±0.67 | 62.45±1.11 | 20.89±1.21 | 8.60±0.20 | 2.88±0.03 | 17.69±1.82 | 6.13±0.37* |
| Mixed-BCDE | 80.56±0.73 | 68.04±1.96 | 61.57±0.32 | 21.31±1.11 | 9.24±0.97 | 2.95±0.06 | 14.99±1.09 | 3.81±0.61 |
| Mono-ABCDE | 79.74±0.27 | 69.33±0.57 | 63.59±1.08 | 20.65±0.97 | 8.59±0.17 | 2.89±0.02 | 17.15±1.48 | 6.13±0.29 |
| Mixed-ABCDE | 80.05±0.62 | 67.64±1.75 | 62.09±0.55 | 21.68±1.13 | 8.83±0.36 | 2.83±0.04 | 15.78±1.15 | 5.78±0.91 |
| Paired t-test |  |  |  |  |  |  |  |  |
| Mono-cropping | 79.83±0.07 | 69.24±0.23 | 63.13±0.56 | 21.52±0.39 | 8.58±0.03 | 2.89±0.01 | 18.66±0.74** | 6.41±0.14** |
| Mixed-cropping | 80.27±0.42 | 69.48±0.45 | 64.75±1.02 | 21.33±0.56 | 9.02±0.21 | 2.89±0.03 | 12.48±0.80 | 4.48±0.37 |

**Table 4.**
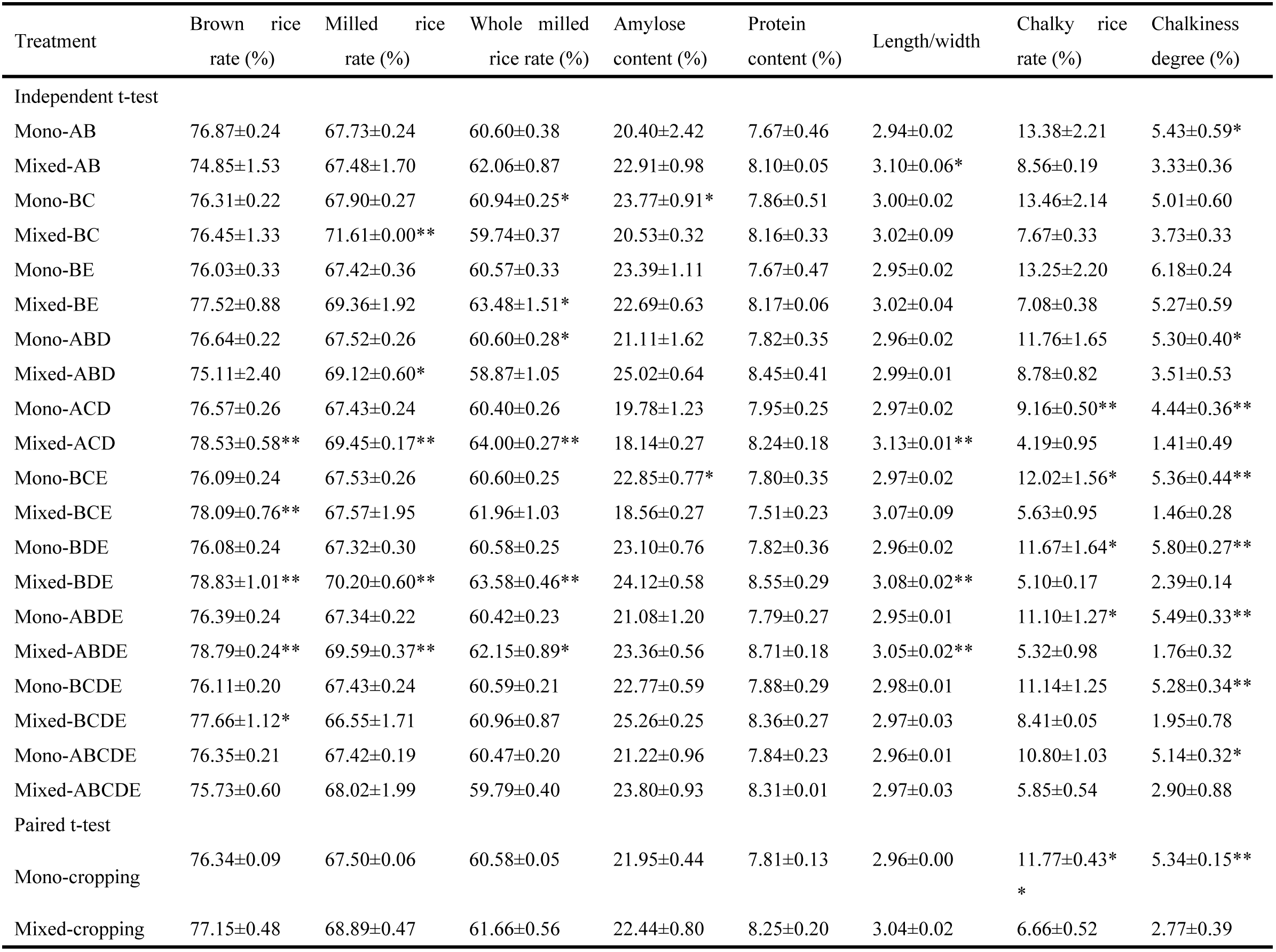
Differences of grain quality between mono-cropping and corresponding mixed-cropping systems with several cultivars in the late growing season of 2016. Note: Table explanations are provided in Table 1.

### Photosynthetic rate, SPAD index and Total Aboveground Dry Weight

In both growing seasons, the photosynthetic rate for all mixed-cropping treatments was higher than that in the mono-cropping treatments, but non-significant for some cases at the maturity stage (Independent t-test, Figs 1a-b). Similar patterns were found for SPAD index and total aboveground dry weight (Independent t-test, Figs 1c-f). Compared with the mono-cropping systems, SPAD index and Pn of the mixed-cropping systems were significantly higher in the early and late growing seasons (pair t-test, Fig. 1). Total aboveground dry weight was also significantly higher for the mixed-cropping treatments in the early growing season but not in the late growing season (pair t-test, Fig. 1). These results indicated that photosynthetic related traits (such as SPAD index and photosynthetic rate) were increased in the mixed-cropping systems, which may in turn enhance aboveground dry weight.

**Fig. 1.**
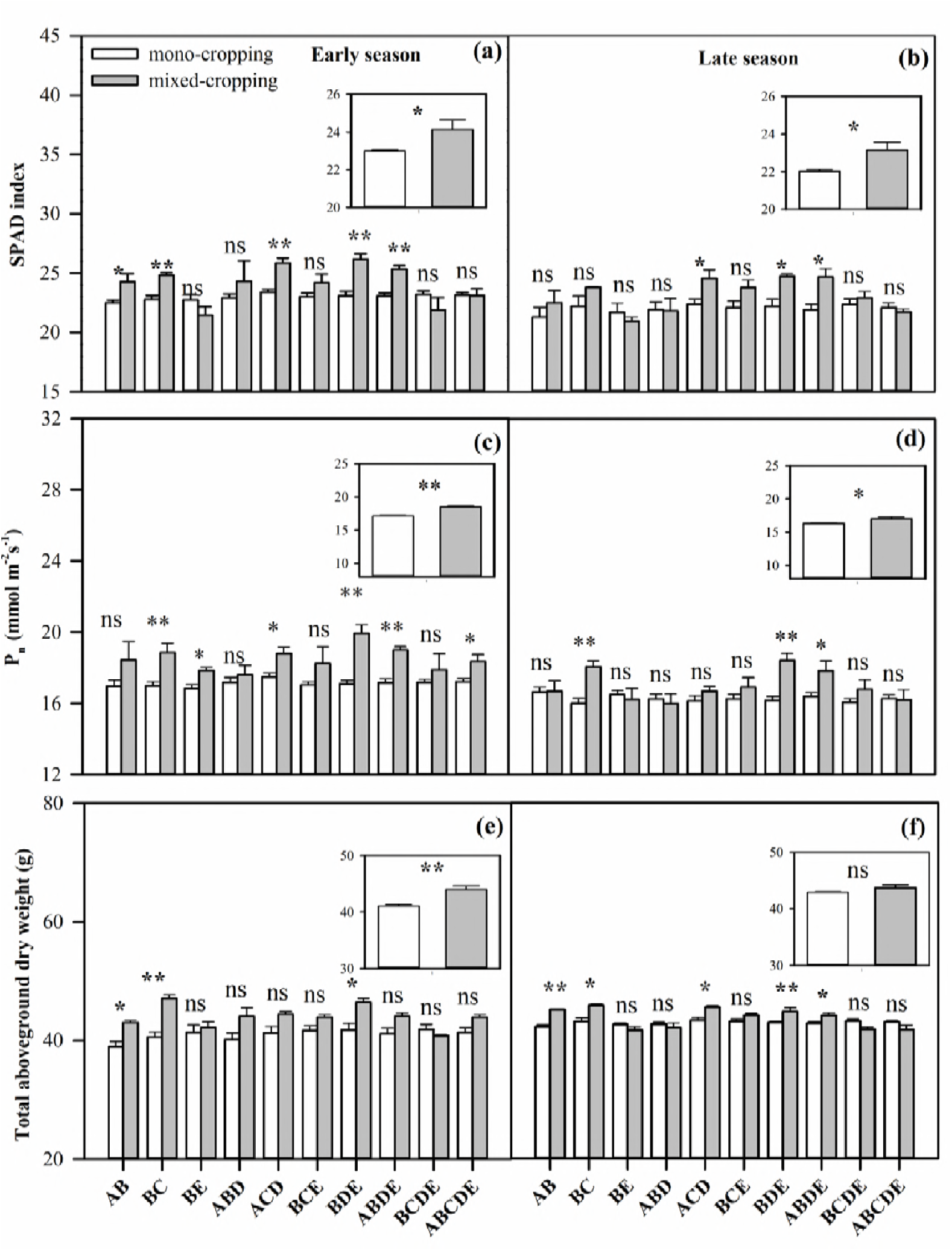
Differences between mono-cropping and corresponding mixed-cropping systems with several cultivars at the maturity stage in the early and late growing season of 2016: (a-b) SPAD index, (c-d) photosynthetic rate (P_n_), and (e-f) total aboveground dry weight. Insets: differences in trait values between mixed-cropping systems and mono-cropping systems (paired t-test). White columns indicate mono-cropping systems and grey columns indicate corresponding mixed-cropping systems.*and**represent significance differences at *P<0.05* and *P<0.01* levels, respectively; ns indicates non-significant.

### Correlations analyses

Pn was significantly (*P< 0.05*) and positively correlated with DW (r=0.67 for the early growing season), seed-setting rate (r=0.59 and r=0.74), grain weight per pot (r=0.53 and r=0.60) and harvest index (r=0.77 and r=0.63) for the early and late growing seasons, respectively (Table 5). A similar pattern was observed for the SPAD index. The results indicated that higher Pn and SPAD index may lead to higher grain yield and yield related traits.

**Table 5.** Correlations analysis between photosynthetic parameters, SPAD index at the maturity stage and yield related traits and grain quality in the early and late growing seasons in 2016.

| Treatments | Early growing season |  | Late growing season |  |
| --- | --- | --- | --- | --- |
|  | SPAD index | Pn | SPAD index | Pn |
| DW | 0.85** | 0.68** | 0.75** | 0.44 |
| Spikelet per Panicle | 0.34 | -0.02 | 0.18 | 0.48 |
| Seed-setting rate | 0.58* | 0.59* | 0.77** | 0.75** |
| Grain weight per pot | 0.74** | 0.54* | 0.88** | 0.60* |
| 1000-grain-weight | 0.65** | 0.29 | 0.45 | 0.58* |
| Harvest index | 0.65** | 0.78** | 0.73** | 0.64* |
| Brown rice rate | 0.28 | 0.56* | 0.55* | 0.64* |
| Milled rice rate | 0.52* | 0.31 | 0.37 | 0.33 |
| Whole milled rice rate | 0.67** | 0.56* | 0.37 | 0.25 |
| Amylose content | -0.29 | 0.09 | -0.11 | -0.23 |
| Protein content | 0.53* | 0.66** | 0.40 | 0.20 |
| Length/width | 0.46 | 0.32 | 0.62* | 0.21 |
| Chalky rice rate | -0.76** | -0.62** | -0.64* | -0.49 |
| Chalkiness degree | -0.40 | -0.63* | -0.65** | -0.38 |

Note: \*and \*\* represent significance at $P < 0.05$ and $0.01$ levels, respectively.

Significantly positive correlations between SPAD index and milled rice rate, whole milled rice rate and protein content were noticed, whilst negative correlations between SPAD index and chalky rice rate were found in the early season (Table 5). Pn was positively correlated with brown rice rate, whole milled rice rate and protein content, whilst negatively correlated with chalky rice rate, chalkiness degree and brown rice rate in the early season. Similar patterns were found in the late season.

## Discussion

Scientists have different opinions regarding mixed-cropping owing to pre-sowing mixing of seeds of different cultivars/genotypes. For instance, some have found this practice was superior to the mono-cropping [24–27], whilst some regarded it as inferior to the mono-cropping regarding yield benefits [28], and others found it depended on the combinations of varieties [29].

Our results showed that compared with mono-cropping systems, mixed-cropping systems indeed have advantages in yield related traits e.g., the number of spikelets per panicle, seed-setting rate, grain weight and harvest index, as well as grain quality traits, e.g. chalky rice rate and chalkiness degree. Positive effects of mixed-cropping systems on plant production generally rely on functional differences between cultivars [30,31]. For example, mixing of seeds of those cultivars having exactly the same functional characteristics would not lead to any additive effect in yield and/or overall production. Previously, it was reported that sometimes seed mixture of different genotypes could lead to yield benefits over individual potential of one cultivar as mono-cropping system. For example, Dubin and Wolfe (1994) [32] reported a 2% increase in grain yield in three-way wheat (*Triticum aestivum* L.) variety mixtures compared with those in pure lines. Helland and Holland (2001) [25] also found an increase of 3% in three oat (*Avena sativa* L.) varieties when grown in mixture rather than mono-crop. Likewise, Gallandt et al. (2001) [33] demonstrated that winter wheat mixtures of two cultivars resulted in 1.5% yield advantage over pure lines, whereas Sarandon and Sarandon (1995) [34] reported that two-way bread wheat mixtures increased the aboveground dry weight by 8% as compared with pure lines.

Our experimental results showed that the Pn, SPAD index and total aboveground dry weight of the mixed-cropping systems were higher than the component cultivars that were grown in pure lines (Fig. 1). Enhanced Pn and SPAD index might lead to the improved performance of rice mixtures. Chlorophylls contents (expressed as SPAD index), is one of the most important factors associated with photosynthetic rate, as well as crop biomass and economic yield in rice [35]. Enhanced photosynthetic rate even at the single-leaf level was recently been found to be a significant contributor in improving crop productivity [36,37]. Furthermore, positive correlations between the photosynthetic rate, SPAD index, total aboveground dry weight and the grain weight per pot in the early and late growing seasons were also observed at the maturity. These results in our study with a real mixed-cropping (growing two or more rice crops simultaneously without definite row arrangement on the same paddy field, i.e. with real mixed-sowing of the seeds in an equal ratio, then mixed-transplanting and finally mixed-harvesting) further corroborate previous study findings which indicate that ‘mixed-cropping systems’ (growing different crop cultivars with definite row arrangements) and ‘intercropping systems’ (growing two or more crops simultaneously with definite row arrangement on the same piece of land) improved the rice stand establishment and resource use efficiency than mono-cropping systems [7,20,38]. In addition, mixed-cropping systems may have benefit in exploiting available resources in a better way than mono-cropping systems and would thus lead to improved grain yield and harvest index [27,29].

Grain quality includes grain appearance and milling, eating, cooking, and nutritional qualities. Genetic, environmental and crop management factors generally affect the grain quality of rice. Our study showed that the chalky rice rate and chalkiness degree of mixed-cropping systems were decreased compared to that of the mono-cropping systems (Tables 3 and 4). Furthermore, negative correlations among SPAD index, Pn and total aboveground dry weight as well as chalky rice rate with chalkiness degree in both growing seasons were recorded. Increased Pn, SPAD index and the total aboveground dry weight observed in the mixed-cropping systems at maturity (Pair t-test, Fig. 1) may in turn contribute to the lower chalky rice rate and chalkiness degree. Several studies found that reduction of Pn at the maturity would increase the occurrence of chalky grains [39,40], while increased Pn of rice leaves resulting in the reduction of chalky rice rate and chalkiness degree.

Furthermore, no significant differences for grain quality traits i.e., brown rice rate, milled rice rate, whole milled rice rate, amylose rate, protein content, or length/width were found between mixed-cropping systems and mono-cropping systems, and also inconsistent for both the growing seasons(Tables 3 and 4). These inconsistent quality parameters during both the growing seasons were possibly related to environmental differences between the early and late growing seasons [41,42]. Effects of cultivating varieties from the same species with mixed seedling, transplanting and harvesting on grain amylose rate, protein content, and length/width are still poorly understood. Previously, in a study focused on a barley-oats mixed system, Jokinen (1991) [43] found that protein content was varied substantially in mixed cropping system rather in mono-cropping system of individual crops.

Overall, the differential crop responses for both mixed-cropping and mono-cropping systems are possibly due to genetic differences among rice cultivars. However, we acknowledge some uncertainties and limitations in our data. For example, firstly, we conducted a pot experiment in this study, whereas pot size, volume, shape, material and even color may influence plant growth [44]. Secondly, we planted rice in a well-controlled greenhouse, and no pests and diseases were detected in both growing seasons, however, the disease resistance of mixed-cropping systems may have more advantages in the field. For example, the effect of mixed cropping systems for pests in organic oilseed crops were evaluated, and the result showed that the infestation by insect pests was directly reduced in mixtures with winter rape (*Brassica napus*) hints and cereals or legumes [45]. Thirdly, with meticulous management and monitoring, there were few weeds in this pot experiment. Indeed, there were some biological effects on weeds-suppressing in mixture with organic linseed (*Linum usitatissivum*) and wheat in the field [45]. Therefore, a field based evaluation of different rice cultivars under mixed cropping system is needed in future.

## Conclusions

In the present study, results showed that the mixed-cropping systems with real mixed-sowing, mixed-transplanting and finally mixed-harvesting have advantages in yield related traits and grain quality compared with the corresponding mono-cropping systems. Relative to the mono-cropping systems, mixed-cropping systems increased the number of spikelets per panicle, seed-setting rate, and grain weight per pot and harvest index. Additionally, mixed-cropping systems reduced chalky rice rate and chalkiness degree. Hence, mixed-cropping system with different rice cultivars would be more productive with higher quality than the mono-cropping systems.

## Acknowledgements

We acknowledge the funding provided by the Science and Technology Project of Guangdong Province (2015B090903077, 2016A020210094), the Science and Technology Project of Guangzhou (201604020062), the Innovation Team Construction Project of Modern Agricultural Industry Technology System of Guangdong Province (2016LM1100), the Overseas Joint Doctoral Training Program of South China Agricultural University (2018LHPY010).

## Supporting information

**S1 Table.** S1 Traits of the five varieties that accessed from the China Rice Data Center. Note: Data from: http://www.ricedata.cn/

| Varieties | Period (d) | Height (cm) | Spikelet per panicle | Seed setting rate (%) | 1000-grain weight (g) | Brown rice rate (%) | Chalky rice (%) | Chalkiness degree (%) | Amylose content (%) | Length/ width | Yield (t hm<sup>2</sup>) |
| --- | --- | --- | --- | --- | --- | --- | --- | --- | --- | --- | --- |
| Yuenongsimiao (A) | 111~113 | 97.0~97.9 | 122~124 | 87.1~88.0 | 22.0~22.6 | 71.8~73.0 | 3~6 | 0.5~0.9 | 17.3~18.2 | 3.3~3.5 | 6.57 |
| Yuxiangyouzhan (B) | 126~128 | 105.6~106.4 | 133.8 | 81.6~86.0 | 22.6 | 46.3~47.0 | 13 | 2.6~8.7 | 3.7~26.3 | / | 6.95 |
| Huangguangyouzhan (C) | 128~132 | 107.7~110.1 | 133~144 | 84.9~87.2 | 24.5~24.6 | 44.0 | 8~11 | 1.0~2.5 | 13.7~15.9 | 3.1 | 7.62 |
| Huanghuazhan (D) | 129~131 | 93.8~102.8 | 118.3~123 | 80.5~86.8 | 22.2~23.1 | 40.0~55.2 | 4~6 | 0.6~3.2 | 13.8~14.0 | / | 7.20 |
| Huahang 31 (E) | 110~111 | 109.5~110.6 | 132.1~132 | 83.5~85.8 | 22~22.3 | 70.4~72.5 | 4~18 | 0.8~6.9 | 16.2~16.5 | / | 6.31 |

